# Endosome-Mediated Epithelial Remodeling Downstream of Hedgehog/Gli Is Required for Tracheoesophageal Separation

**DOI:** 10.1101/777664

**Authors:** Talia Nasr, Pamela Mancini, Scott A. Rankin, Nicole A. Edwards, Zachary N. Agricola, Alan P. Kenny, Jessica L. Kinney, Keziah Daniels, Jon Vardanyan, Lu Han, Stephen L. Trisno, Sang-Wook Cha, James M. Wells, Matthew J. Kofron, Aaron M. Zorn

**Author notes:** These authors contributed equally (Talia Nasr, Pamela Mancini).

## Abstract

The trachea and esophagus arise from the separation of a common foregut tube during early fetal development. Mutations in key signaling pathways such as Hedgehog (HH)/Gli can disrupt tracheoesophageal (TE) morphogenesis and cause life-threatening birth defects (TEDs), however the underlying cellular mechanisms are unknown. Here we use mouse and *Xenopus* to define the HH/Gli-dependent processes orchestrating TE morphogenesis. We show that downstream of Gli the Foxf1+ splanchnic mesenchyme promotes medial constriction of the foregut at the boundary between the presumptive Sox2+ esophageal and Nkx2-1+ tracheal epithelium. We identify a unique boundary epithelium co-expressing Sox2 and Nkx2-1 that fuses to form a transient septum. Septum formation and resolution into distinct trachea and esophagus requires endosome-mediated epithelial remodeling involving the small GTPase Rab11, and localized extracellular matrix degradation. These are disrupted in Gli-deficient embryos. This work provides a new mechanistic framework for TE morphogenesis and informs the cellular basis of human TEDs.

**Highlight bullet points:**

- The Sox2+ esophagus and Nkx2-1+ trachea arise from the separation of a single foregut tube through a series of cellular events conserved in mouse and *Xenopus*
- Tracheoesophageal morphogenesis initiates with HH/Gli-dependent medial constriction of the gut tube mesenchyme at the Sox2-Nkx2-1 border
- The foregut epithelial walls fuse forming a transient septum co-expressing Sox2 and Nkx2-1
- Downstream of HH/Gli Rab11-dependent endosome-mediated epithelial remodeling and localized extracellular matrix degradation separate the esophagus and trachea
- HH/Gli mutations reveal the cellular basis of tracheoesophageal birth defects

## INTRODUCTION

Between 25-35 days of human gestation, the fetal gut tube separates into the distinct trachea and esophagus (Billmyre et al., 2015; Que, 2015). Disruptions in this process result in life-threatening defects that impair neonatal respiration and feeding, including esophageal atresia (EA), tracheal atresia (TA), and/or tracheoesophageal fistula (TEF) (Brosens et al., 2014). The etiology of TEDs, occurring in ∼1:3500 births, is poorly understood (Shaw-Smith, 2006). Even when patient genetics or animal models have revealed the genes involved, how mutations result in TEDs is unclear because the cellular basis of TE morphogenesis are poorly defined.

Studies in mouse and *Xenopus* have shown that TE development is initiated by a signaling cascade involving HH, WNT and other signals between the foregut endoderm and the surrounding splanchnic mesoderm. These signals pattern the epithelium into a ventral respiratory domain expressing the transcription factor Nkx2-1 and a dorsal esophageal domain expressing the transcription factor Sox2 around mouse embryonic day (E) 9.0 and *Xenopus* NF35, 50 hours post fertilization (hpf) (Hines and Sun, 2014; Rankin et al., 2016). Over the next few days, the foregut separates into distinct TE tubes, with paired lung buds emerging from the posterior aspect of the Nkx2-1+ domain via an Fgf10-mediated mechanism that is distinct from TE separation (Hines and Sun, 2014).

Mouse mutations in these patterning genes can result in TEDs similar to those seen in patients. For example, *Sox2* and *Nkx2-1* knockouts result in EA and TA, respectively (Minoo et al., 1999; Que et al., 2007; Trisno et al., 2018), while HH pathway mutations, such as in the ligand *Shh* or the transcription factor*s Gli2* and *Gli3*, can cause a spectrum of defects ranging from EA/TEF to laryngotracheoesophageal clefts (LTECs) (Litingtung et al., 1998; Motoyama et al., 1998; Rankin et al., 2016; Tabler et al., 2017). How these mutations result in TEDs is unclear since the cellular processes that HH regulates in this context are unknown. While a number of models for TE morphogenesis have been postulated (Billmyre et al., 2015; Que, 2015), the underlying cellular mechanisms remain to be elucidated.

Here we use *Xenopus* and mouse to define the conserved cellular mechanisms orchestrating TE morphogenesis. We show that HH/Gli signaling regulates multiple steps of TE separation, including dorsal-ventral (D-V) patterning, medial constriction and endosome-mediated epithelial remodeling. These results provide a mechanistic foundation for TE morphogenesis to inform the genotype-phenotype basis of human TEDs.

## RESULTS and DISCUSSION

### TE Morphogenesis is Conserved in *Xenopus* and Mouse

To define the cell biology of TE morphogenesis, we took a comparative approach using mouse and *Xenopus*. TE development appears to be conserved with patterning of the foregut epithelium into Sox2+ dorsal and Nkx2-1+ ventral domains by NF35 in *Xenopus* and E9.5 in mouse. The foregut then separates into distinct TE tubes by NF42 (80 hpf) in *Xenopus* and E11.5 in mice (Figures 1A-C’ and S1A-F’) (Rankin et al., 2015). Close examination of *Xenopus* embryos over the course of TE morphogenesis allowed us to classify the process into four major steps: 1) D-V patterning; 2) medial constriction at the Sox2-Nkx2-1 boundary; 3) epithelial fusion and remodeling of a transient septum; and 4) mesenchymal invasion separating the TE tubes. We investigated the cellular basis of each step, first using *Xenopus* where we could screen many potential mechanisms, followed by a comparison to mouse.

**Figure 1:**
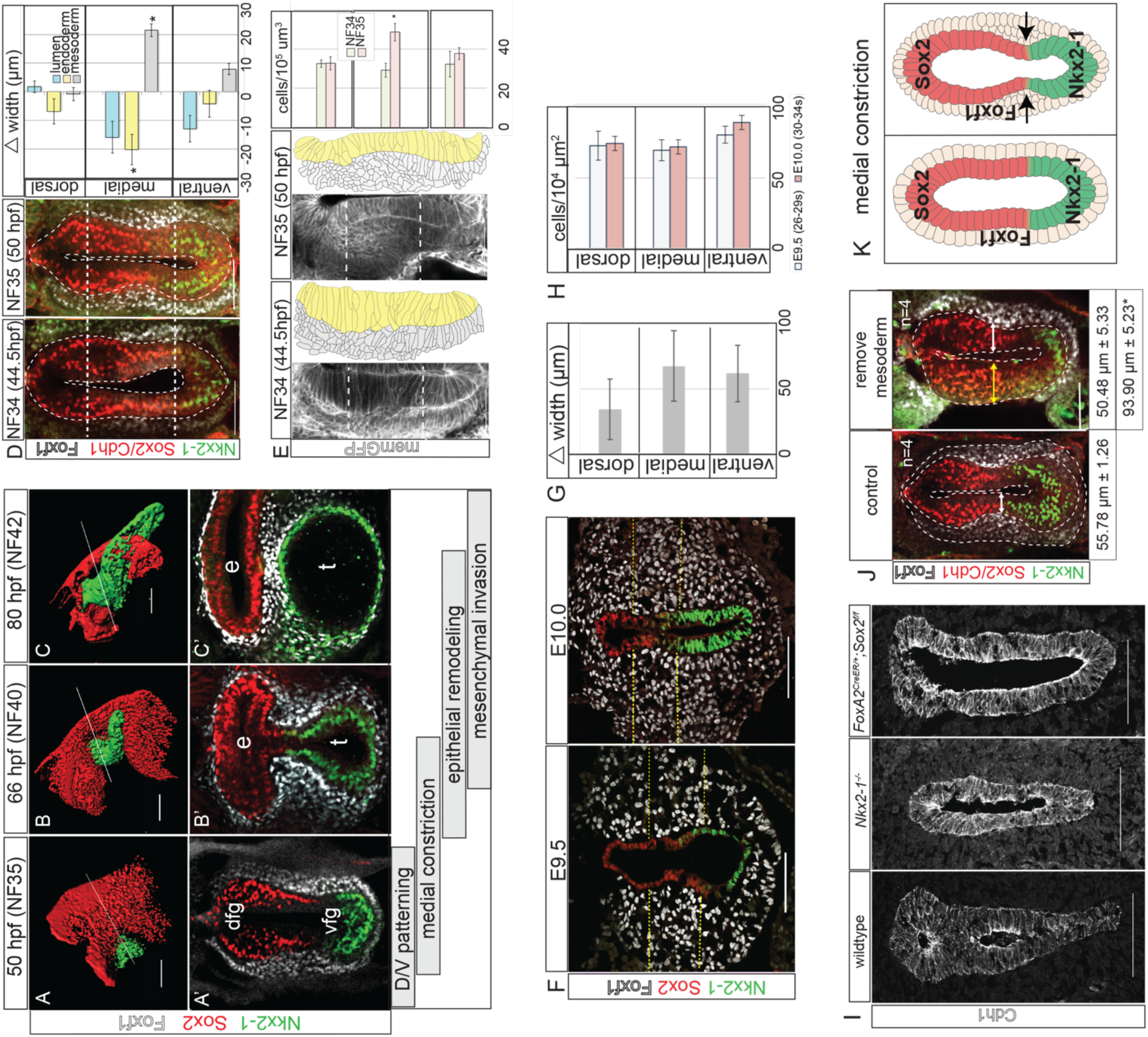
Foxf1+ Mesenchyme Promotes Medial Constriction at the Sox2-Nkx2-1 Boundary. 1A-C’: Immunostaining showing TE morphogenesis in *Xenopus (X*.) *laevis.* Scale bar, 100 μm. 1D: Medial constriction in *X. laevis* shown by immunostaining and quantification of the difference in foregut width between NF34 and NF35. Scale bar, 100 μm. Difference of means test,*p<0.5. 1E: Transgenic membrane-GFP *X. laevis* show increased medial mesenchyme cell density. Scale bar, 100 μm. Student’s two-tailed t-test,*p<0.05. 1F: Immunostaining of mouse embryos showing medial constriction. Foxf1+ cells between dashed yellow arrows denote medial mesoderm. Scale bar, 100 μm. 1G: Average mesoderm width was not significantly different between E9.5 and E10.0 in the dorsal, medial, and ventral regions. Difference of means test. 1H. Average mesoderm cell density at E9.5 and E10.0 was not significantly different. Student’s two-tailed t-test. 1I: E11.0 *Nkx2-1* and E10.5 *Sox2* mouse mutants fail to undergo medial constriction. Scale bar, 100 μm. 1J: Removal of the lateral plate mesoderm prevents medial constriction in *X. laevis* embryos. Student’s two-tailed t-test performed between control and explanted widths, and explanted and intact widths.*p<0.05. Scale bar, 100 μm. 1I: Graphical summary of medial constriction.

### Foxf1+ Mesenchyme Promotes Medial Constriction at the Sox2-Nkx2-1 Boundary

In *Xenopus*, the first morphological indication of TE separation was a medial constriction of the foregut at the Sox2-Nkx2-1 boundary by NF35, with a significant narrowing of the gut tube lumen coincident with thickening of the medial Foxf1+ splanchnic mesoderm (Figure 1D). Analysis of transgenic membrane-GFP embryos revealed that the mesoderm thickening was not due to increased mesodermal cell size or cell shape changes, but rather to a condensation of Foxf1+ mesodermal cells (Figures 1E and S1K). This was accompanied by the epithelium transitioning from a thick pseudostratified layer at NF34 to a thinner columnar epithelium at NF35 (Figure 1E). Phospho-Histone H3 staining indicated that the medial mesenchyme had a slightly higher proliferation rate than the dorsal or ventral regions, which may in part explain the localized increase in mesenchymal cell numbers (Figure S1G,H).

The mouse foregut also constricted at the Sox2-Nkx2-1 boundary between E9.5 and E10.0 (Figure 1F), but there was no relative increase in medial mesoderm thickness, cell density or proliferation compared to the ventral or dorsal tissue as observed in *Xenopus* (Figures 1G, H and S1I-L). In general, the mesoderm in mouse is much thicker than in *Xenopus*; however, in both species the Foxf1+ cells were more tightly packed against the epithelium at the constriction point (Figures 1A-F and S1I), suggesting a close association of the mesenchyme with the basement membrane in this region.

Since foregut constriction occurs at the Sox2-Nkx2-1 boundary, we re-examined mouse *Nkx2-1* and *Sox2* mutants where the TE does not separate, but the underlying mechanisms have not been reported. Both *Nkx2-1* and *Sox2* mutants failed to undergo medial constriction (Figure 1I), indicating that proper D-V patterning and the function of these two lineage-determining transcription factors are essential to initiate TE morphogenesis. We postulate that the Sox2-Nkx2-1 boundary might also have some inherent information that regulates local mesenchyme behavior. These data help explain why human *SOX2* mutations frequently cause EA; if the SOX2+ domain is too small, foregut constriction may not occur or might initiate too dorsally, resulting in insufficient esophageal tissue.

Since mesenchymal condensations can promote bending of adjacent epithelial sheets in some contexts (Walton et al., 2012), we tested whether the mesoderm was required for medial constriction of the *Xenopus* gut tube by surgically removing the splanchnic mesenchyme from one side of the NF32 foregut. Imaging at NF35 revealed a failure of medial constriction and epithelial thinning relative to the contralateral control side (Figure 1J), suggesting that the mesoderm is required for medial constriction, possibly by exerting a pushing force on the epithelium. Together, these data indicate that medial constriction of the foregut at the Sox2-Nkx2-1 boundary is the critical first step in TE morphogenesis (Figure 1K) and that disruptions in this process can lead to TEDs.

### Rab11 and Endosome-Mediated Epithelial Remodeling are Required for TE Septation

As the foregut constricts, the opposing epithelial walls come into contact, forming a transient epithelial septum (Figures 2A-C). Close examination revealed a unique cell population in the septum that co-expresses Nkx2-1 and Sox2 (Figures 2D and S2A). We examined whether modulators of epithelial behavior were differentially active at the Sox2+/Nkx2-1+ contact point. Before collapse of the medial lumen, the endoderm is a polarized epithelium with aPKC on the apical/luminal surface and Cdh1 enriched in adherens junctions and on the basolateral surface. As the epithelial walls touched, aPKC was specifically reduced at the contact point (Figure 2B-C), where the transient septum formed like a closing zipper between the presumptive esophagus and trachea. Concurrently, Cdh1 and Integrin became enriched on what was previously the apical surface (Figure 2B-F), suggesting that cell-cell adhesion holds the two sides together.

**Figure 2:**
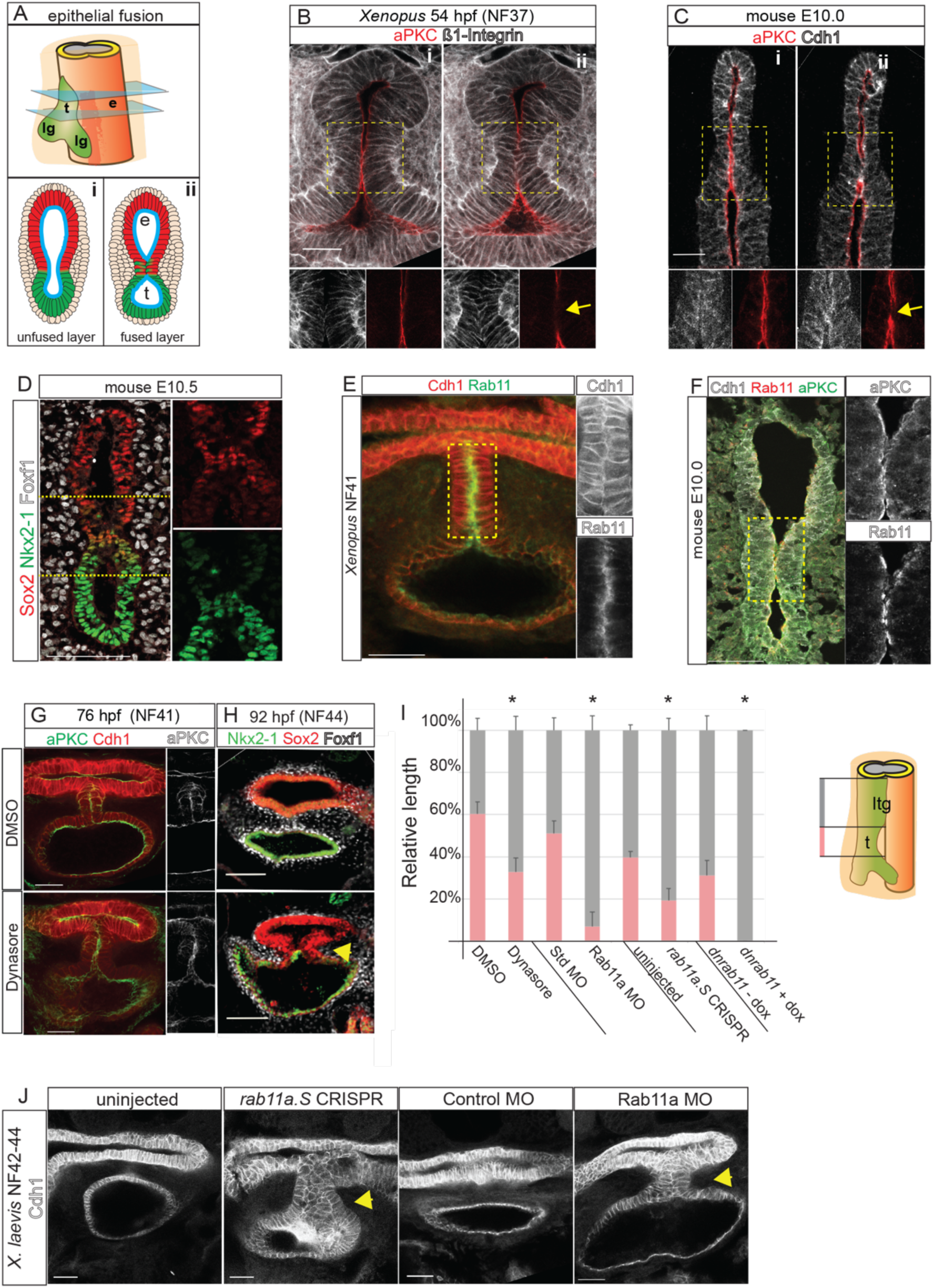
Endosome-Mediated Epithelial Remodeling is Required for TE Septation. 2A: Model of epithelial fusion. 2B-C: Sequential optical sections of *X. laevis* (B) and mouse (C) embryo immunostaining showing loss of aPKC and increased Integrin or Cdh1 at the contact point (arrow, ii). Scale bar, 50 μm. 2D: A unique population of cells co-expressing of Sox2 and Nkx2-1 in the mouse foregut septum. Scale bar, 100 μm. 2E: Immunostaining of Rab11 and Cdh1 enriched in the *X. laevis* septum. Scale bar, 50 μm. 2F: Immunostaining of Rab11, aPKC and Cdh1 enriched at the fusion point in mouse. Scale bar, 50 μm. 2G-H: Inhibition of endosome recycling by Dynasore treatment of *X. laevis* (NF32-41) results in a failure to reduce aPKC at NF41 (G) and tracheoesophageal cleft (TEC) at NF44 (H). Scale bar, 100 μm. 2I: Quantification of reduced trachea (t) length and relative to laryngotracheal (ltg) in NF42-44 *X. laevis* embryos shows that inhibition of Rab11-mediated endocytosis results in a TEC. Student’s two-tailed t-test, *p<0.05. 2J. Cdh1 immunostaining of NF42-44 *X. laevis* showing that knockdown of Rab11a via CRISPR-mediated mutation or MO knockdown results in a TEC. Scale bar, 50 µm.

Remodeling of polarized epithelium is regulated by endosome recycling, where aPKC and cadherins are removed from the cell surface by endocytosis and shuttled to other membrane domains (Ivanov et al., 2004; Wang et al., 2018). Immunostaining for endosomal protein Rab11 revealed abundant apical puncta where the epithelium fuses in both *Xenopus* and in mouse, coincident with loss of aPKC and Cdh1 relocalization (Figure 2E-F). To test whether endosome recycling is required for TE separation, we treated *Xenopus* embryos at NF32 with Dynasore, a dynamin inhibitor that blocks endocytosis (Macia et al., 2006). In Dynasore-treated embryos, aPKC remained aberrantly localized on the apical surface of the epithelial contact site at NF41, resulting in a tracheoesophageal cleft (TEC) at NF44 (Figure 2G-I).

To corroborate the pharmacological inhibition, we performed F0 CRISPR/Cas9-mediated indel mutation of *rab11a* in *Xenopus* (Methods), as well as Rab11a knockdown with a well-characterized Rab11a antisense morpholino oligo (MO) (Kim et al., 2012). All of these resulted in 25-30% of the embryos having gastrulation defects, as expected (Kim et al., 2012; Ossipova et al., 2015); however ∼50% of the remaining embryos (which were grossly normal) exhibited a TEC at NF42-43 with a failure of the transient epithelial septum to resolve (Figures 2I,J and S2E). In the case of *rab11a*-CRISPRs and Rab11a-MO, immunostaining confirmed the reduced Rab11 protein (Figure S2B). Finally, to overcome the earlier role of Rab11 in gastrulation, we generated transgenic *Xenopus* embryos expressing a dominant negative Rab11 in the foregut upon induction with doxycycline [*Tg(hhex:trTA;TRE:dnRab11a-GFP)*]. These also resulted in a TEC with enrichment of the dnRab11a-GFP on the basal-lateral epithelium rather than the normal apical surface (Figure S2C-E). Together, these data demonstrate that Rab11a-dependent endosome-mediated epithelial remodeling is essential for the foregut walls to fuse properly and that disruptions in this process can lead to TEDs.

### Localized ECM Degradation and Mesenchymal Invasion Resolve the TE Septum

Between NF40-42 in *Xenopus* and E10.5-11.5 in mice, the length of the Sox2-Nkx2-1 boundary decreased as the length of the separated trachea and esophagus increased (Figure 3A). The presumptive trachea and esophagus had comparable proliferation in both species (data not shown), indicating that differential growth is unlikely to account for TE separation (Ioannides et al., 2010). This is consistent with a “splitting and extension” model (Que, 2015), where active septation occurs along with relatively equal growth of the separated trachea and esophagus.

**Figure 3:**
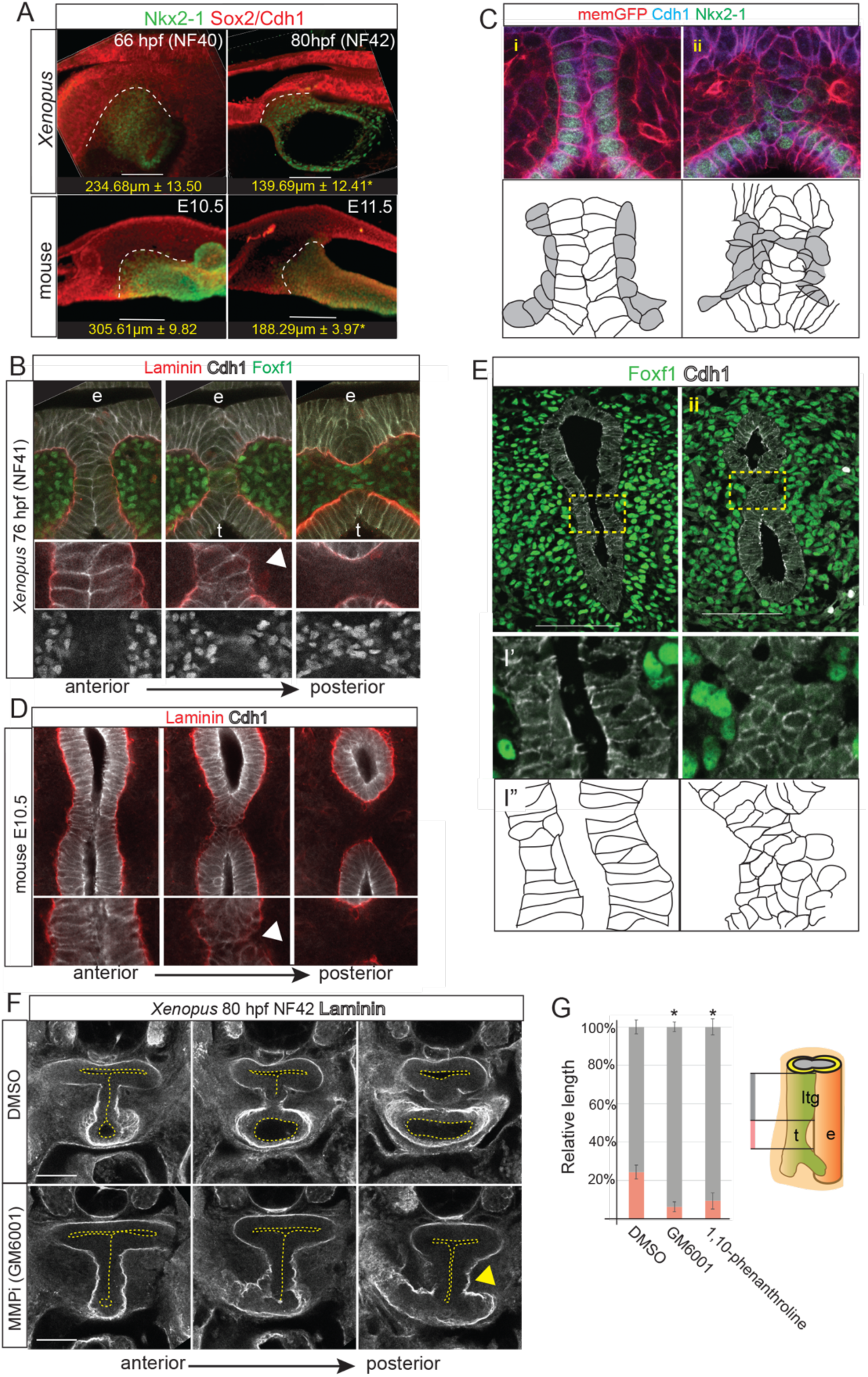
Localized ECM Degradation and Mesenchymal Invasion Resolve the TE Septum. 3A: Wholemount immunostaining of septum resolution in *X. laevis* and mouse embryos, quantifying the length of the Sox2+/Nkx2-1+ boundary. Student’s two-tailed t-test,*p<0.05. 3B: Optical sections from wholemount immunostaining of Laminin, Cdh1 and Foxf1 during TE septum resolution in NF41 *X. laevis* embryos. Arrowhead indicates localized Laminin breakdown. 3C: Immunostaining of transgenic membrane GFP NF41 *Xenopus* embryo before (i) and during (ii) TE separation showing Cdh1+ epithelial cells (white in schematic) round up as mesenchymal cells (grey) invade. Scale bar, 100 µm. 3D: Optical sections from wholemount immunostaining of Laminin and Cdh1 during TE septum resolution in E10.5 mouse embryos. Arrowhead points to localized Laminin breakdown. 3E: Immunostaining of Foxf1 and Cdh1 (schematic below) in an E10.5 mouse embryo before (i) and during (ii) TE separation. Scale bar, 100 μm. 3F: Inhibition of MMP activity in *Xenopus* with GM6001 results in impaired Laminin breakdown (arrowhead) and result in a TEC. Three optical sections from wholemount Laminin immunostaining of NF41 *X. laevis* embryos. 3G: Quantification of relative lengths of the laryngotracheal groove (ltg) and trachea (t) in DMSO, GM6001, or 1,10-phenanthroline-treated NF41 *X. laevis* embryos. One-way ANOVA, *p<0.05.

3D confocal wholemount staining indicate that septation occurs in a posterior to anterior wave starting at the lung buds (Figures 3A-E, S3 and Supplementary Movies). In both *Xenopus* and mouse, the anterior septum was composed of Cdh1+ columnar cells and Laminin-rich basement membrane surrounded the entire foregut epithelium. However, in more posterior optical sections, Cdh1 and Laminin levels became reduced as basement membrane broke down and epithelial cells in the septum rounded up and lost adhesion to one another (Figure 3B-E and Supplementary Movies). Cdh1+ puncta indicative of new adherens junctions were observed between opposing epithelial cells to seal the presumptive tracheal and esophageal lumens (Figures 3B-D). Inhibiting ECM remodeling matrix metalloproteinases (MMPs) (Vu and Werb, 2000) with either GM6001 or 1,10-phenanthroline impaired Laminin breakdown in *Xenopus*, resulting in a TEC and shorter trachea (Figure 3F-G). As the septum resolved, fibronectin (Fn1)-rich Foxf1+ mesoderm cells, with enriched cortical actin, were observed between the separating trachea and esophagus, suggesting that mesenchymal cells actively migrate across the midline on the Fn1+ matrix (Figure S3C, D and F).

We evaluated the possibility that the septum cells undergo apoptosis. Cleaved caspase-3 staining revealed a few dying cells in the resolving mouse septum (Figure S3G) (Ioannides et al., 2010), but little septum cell death in *Xenopus* (data not shown). Rather, when the septum cells lost adhesion to one another, they appeared to incorporate into the esophageal or tracheal epithelium. This is consistent with reports in the companion paper (Kim et. al, 2019) and our observation that, immediately after TE separation in both *Xenopus* and mouse, it is not uncommon to see Nkx2-1+ cells in the ventral esophagus and Sox2+ cells in the dorsal trachea (Figure S1E). Thus, in both *Xenopus* and mouse, the transient septum is resolved by epithelial remodeling, localized ECM degradation, and mesenchymal cell invasion to separate the foregut into a distinct esophagus and trachea (Figure S3H). Disruption of this process at any point along the anterior-posterior axis could result in a TEF.

### HH/Gli Activity is Required for D-V Patterning, Medial Constriction and Epithelial Remodeling

Having defined the cellular events controlling TE morphogenesis, we next asked how disruptions in the HH signaling pathway cause TEDs. HH ligands are expressed in the foregut epithelium and signal to the mesoderm to activate Gli2 and Gli3 (Ioannides et al., 2003; Rankin et al., 2016). In the absence of a HH signal, Gli2 is degraded and Gli3 is proteolytically cleaved into a transcriptional repressor (Gli3R). When the HH pathway is activated, Gli2 and Gli3 become transcriptional activators (Briscoe and Therond, 2013). Multiple human TEDs have been associated with mutations in *SHH, GLI2*, and *GLI3*, including heterozygous truncating mutations in *GLI3* (Johnston et al., 2005).

We examined an allelic series of HH/Gli mouse mutants to determine which steps in TE morphogenesis were disrupted. Consistent with previous reports (Litingtung et al., 1998), *Shh*^-/-^ embryos exhibited TA with a single hypomorphic gut tube at E15.5 (Figure 4A). *Gli2*^*-/-*^;*Gli3*^*+/-*^ mutants also failed to undergo TE separation, displaying a TEC along the length of the foregut (Figure 4A) (Motoyama et al., 1998). In contrast, *Gli2*^*+/-*^;*Gli3*^*-/-*^, *Gli3*^*-/-*^ and *Gli2*^*-/-*^ (Figure 4A and data not shown) mutants were indistinguishable from wildtype littermates, indicating a qualitative difference in Gli2 and Gli3, with a single copy of Gli2 being sufficient to support TE morphogenesis. We also examined transgenic embryos ectopically expressing a truncated repressor form of Gli3 (Vokes et al., 2008) in the foregut mesendoderm (*Foxg1Cre;Gli3T*), which mimics the *GLI3* mutation in Pallister-Hall Syndrome (PHS) [Online Mendelian Inheritance in Man 151560] (Johnston et al., 2005). Like PHS patients, *Foxg1;Gli3T* embryos had LTECs (Figure 4A). These results indicate that increasing levels of Gli3R relative to Gli activator cause increasing severity of TEDs.

**Figure 4:**
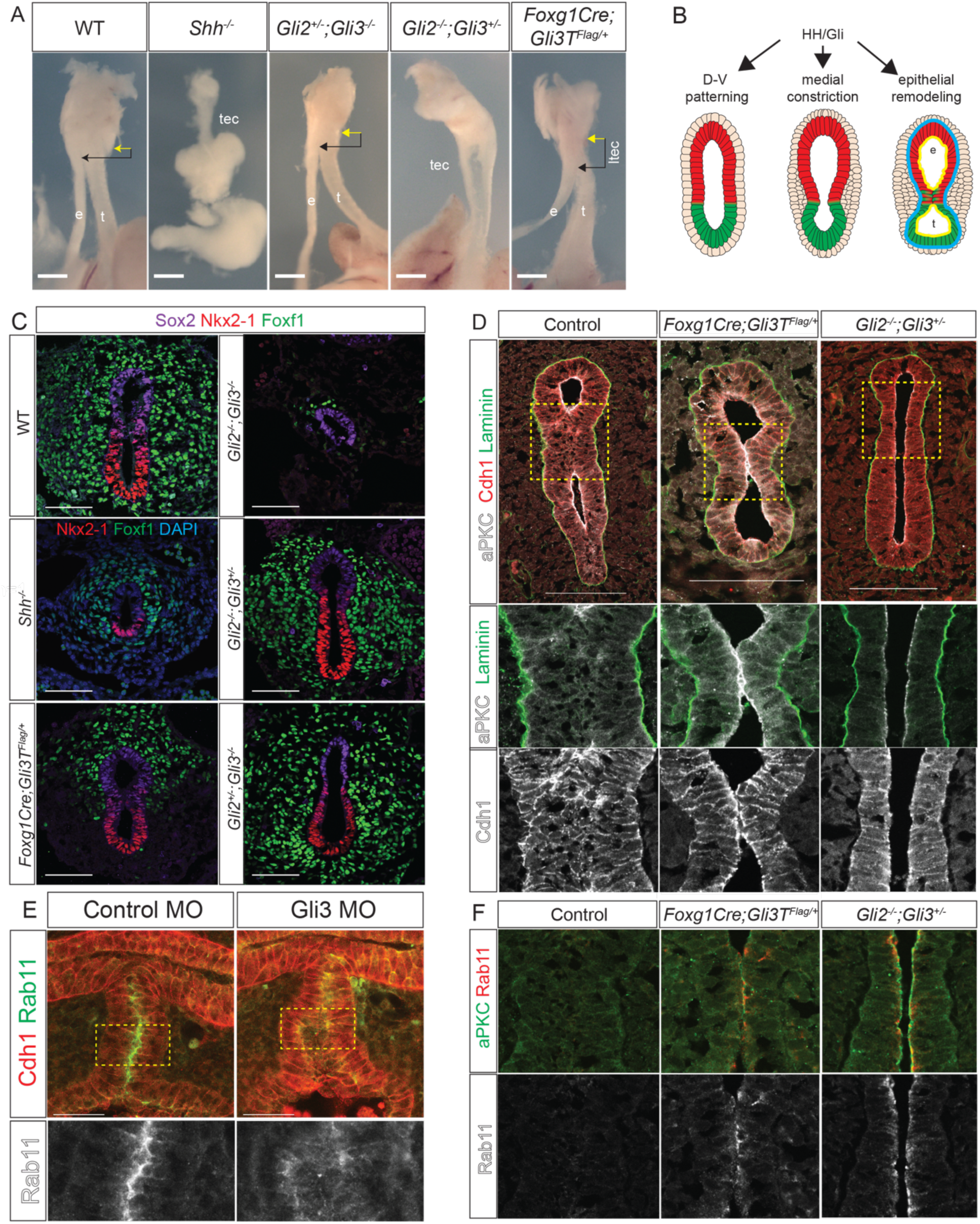
HH/Gli Activity Is Required for D-V Patterning, Medial Constriction and Epithelial Remodeling. 4A: Wholemount images of E15.5 *Shh/Gli* mutant foreguts showing a separate esophagus (e) and trachea (t); tracheoesophageal cleft (tec), or LTEC. Arrows denote distance between cricoid cartilage (yellow) and TE septation point (black). Scale bar, 6.35 mm. 4B: Graphical summary of HH/Gli-regulated events. 4C: Immunostaining of Nkx2-1, Foxf1 and Sox2 (or DAPI) in *Shh*/*Gli* mutants at E10.0. Scale bar, 50 µm. 4D: Immunostaining of aPKC, Laminin and Cdh1 in *Foxg1Cre;Gli3T* and *Gli2*^*-/-*^;*Gli3*^*+/-*^ E11.0 mutants showing a failure epithelial fusion and persistent aPKC. Scale bar, 100 µm. 4E: Immunostaining of control MO and Gli3 MO injected NF41 *X. laevis* embryos showing mislocalized Rab11 in Gli3 morphants. Scale bar, 50 µm. 4F: Immunostaining of aPKC and Rab11 in *Foxg1Cre;Gli3T* and *Gli2*^*-/-*^;*Gli3*^*+/-*^ E11.0 mutants showing a failure of Rab11 reduction compared to controls.

To understand the cellular basis of these defects (Figure 4B), we examined earlier developmental stages. At E10.0, *Gli2*^*-/-*^;*Gli3*^*-/-*^ embryos, which lack all HH activity, failed to undergo D-V patterning, with no Nkx2-1+ respiratory progenitors and very little Foxf1+ mesenchyme (Rankin et al., 2016) (Figure 4C). In contrast, the foregut was properly patterned in all other genotypes with ventral Nkx2-1 and dorsal Sox2, although the *Shh*^*-/-*^ foregut was smaller with fewer Nkx2-1+ and Foxf1+ cells. Although *Shh*^-/-^ mutants were correctly patterned, they failed to medially constrict, similar to NF37 *Xenopus* embryos treated with HH antagonist cyclopamine (Figures 4C and S4B). Unlike *Shh*^*-/-*^ mutants, *Gli2*^*-/-*^;*Gli3*^*+/-*^ embryos had normal levels of Foxf1+ mesenchyme and initiated medial constriction. *Foxg1Cre;Gli3T* embryos also exhibited medial constriction, although these had fewer Foxf1+ cells in the ventral region (Figure 4C). These data show that high levels of HH activity are needed for D-V patterning, medial constriction and to maintain sufficient Foxf1+ mesoderm, consistent with *Foxf1* being a direct Gli transcriptional target (Mahlapuu et al., 2001).

We next assessed *Gli2*^*-/-*^;*Gli3*^*+/-*^ and *Foxg1Cre;Gli3T* mutants at E11.0 for defects in epithelial remodeling and septation (Figure 4D). In *Gli2*^*-/-*^;*Gli3*^*+/-*^ mutants the foregut lumen remained open, suggesting that although medial constriction was initiated, the process was insufficient to bring the epithelial walls into contact. Alternatively, disrupted epithelial remodeling might result in failure of cell adhesion at the constriction point, followed by a relaxation and a TEC. Indeed, in *Foxg1Cre;Gli3T* embryos, aPKC and Rab11 persisted at the anterior contact point where the epithelium touched but failed to fuse, whereas in wild type embryos aPKC, Rab11 and Cdh1 were rapidly downregulated as the opposite walls of the epithelium fused (Figure 4F). This was similar to *Xenopus* embryos with disrupted endosome recycling, suggesting that the LTEC was caused by impaired epithelial remodeling in the septum.

Gli3 MO knockdown in *X. laevis* and CRISPR/Cas9-mediated *gli3* mutation in *X. tropicalis* resulted in phenotypes similar to *Gli2*^*-/-*^;*Gli3*^*+/-*^ and *Foxg1Cre;Gli3T* mouse mutants, with a delay in medial constriction and failure of TE separation resulting in TECs (Figure S4). In most cases, the transient septum formed but failed to resolve, with the lumen eventually reopening to form a cleft. CRISPR-mediated indel mutations of the *gli3* C-terminus, predicted to result in a Gli3R form like the PHS patient mutation, resulted in persistent aPKC in the septum, like *Foxg1Cre;Gli3T* mouse embryos. Immunostaining of Gli3 MO *X. laevis* embryos showed a mislocalization of Rab11, which is normally enriched at the point of apical membrane fusion (Figure 4E). These results collectively indicate that HH/Gli-regulated epithelial remodeling is required for TE morphogenesis in both *Xenopus* and mouse, and that this is compromised in *Gli3* mutant models of PHS.

### Conclusion

We have defined the conserved HH/Gli-regulated cellular mechanisms orchestrating TE morphogenesis: 1) D-V foregut patterning; 2) mesenchymal medial constriction at the Sox2-Nkx2-1 boundary; 3) epithelial fusion of cells co-expressing Sox2 and Nkx2-1 to form a transient septum; and 4) Rab11-dependent endosome-mediated epithelial remodeling and localized ECM breakdown to resolve the septum. Importantly, these results provide a cellular explanation for the previous models of TE separation. The data also help explain the genotype-phenotype association of TEDs in patients with *SHH* and *GLI* mutations, as a greater loss of HH activity causes earlier and more severe TE morphogenesis disruptions. Complete loss of HH/Gli results in TA, while one copy of Gli3 appears to be sufficient to support D-V patterning and constriction, but not septation, resulting in a TEC. We predict that partial or transient disruption in epithelial remodeling would result in an interruption in progressive septation and a TEF. Our data also suggest that a balance between Gli activator and repressor function is critical, with excessive Gli3R resulting in TEDs. Further studies will focus on identifying mesenchymal Gli targets and how these regulate epithelial remodeling and TE morphogenesis. Candidates include proteins that modulate cell adhesion, endosome-mediated epithelial remodeling, and ECM degradation. Consistent with this prediction, human TEDs have been linked to mutations in Integrin Alpha 6 (*ITGA6*), Filamin A (*FLNA*), and Fraser extracellular matrix subunit 1 (*FRAS1)* (Brosens et al., 2014).

## ACKNOWLEDGMENTS

The authors would like to thank the Zorn and Wells labs for their feedback. We acknowledge the CCHMC Confocal Imaging Core. This project was funded by NIH P01HD093363 and was supported in part by NIH P30 DL0778392 to the Digestive Diseases Research Core Center in Cincinnati. T.N. is supported by NIH F30HL142201 and T32 GM063483-14 to the University of Cincinnati Medical Scientist Training Program. We are grateful to Sergei Sokol for the dnRab11 constructs.

## AUTHOR CONTRIBUTIONS

T.N. and A.M.Z. wrote the manuscript. T.N., P.M., and A.M.Z. designed and performed experiments. S.A.R., N.A.E., Z.N.A., A.P.K., J.L.K., K.D., J.V., L.H., S.L.T., M.J.K., S.C., and performed experiments. J.M.W. provided critical discussion and support. All authors provided feedback on the manuscript.

## SUPPLEMENTAL FIGURES

**Supplemental Figure S1.**
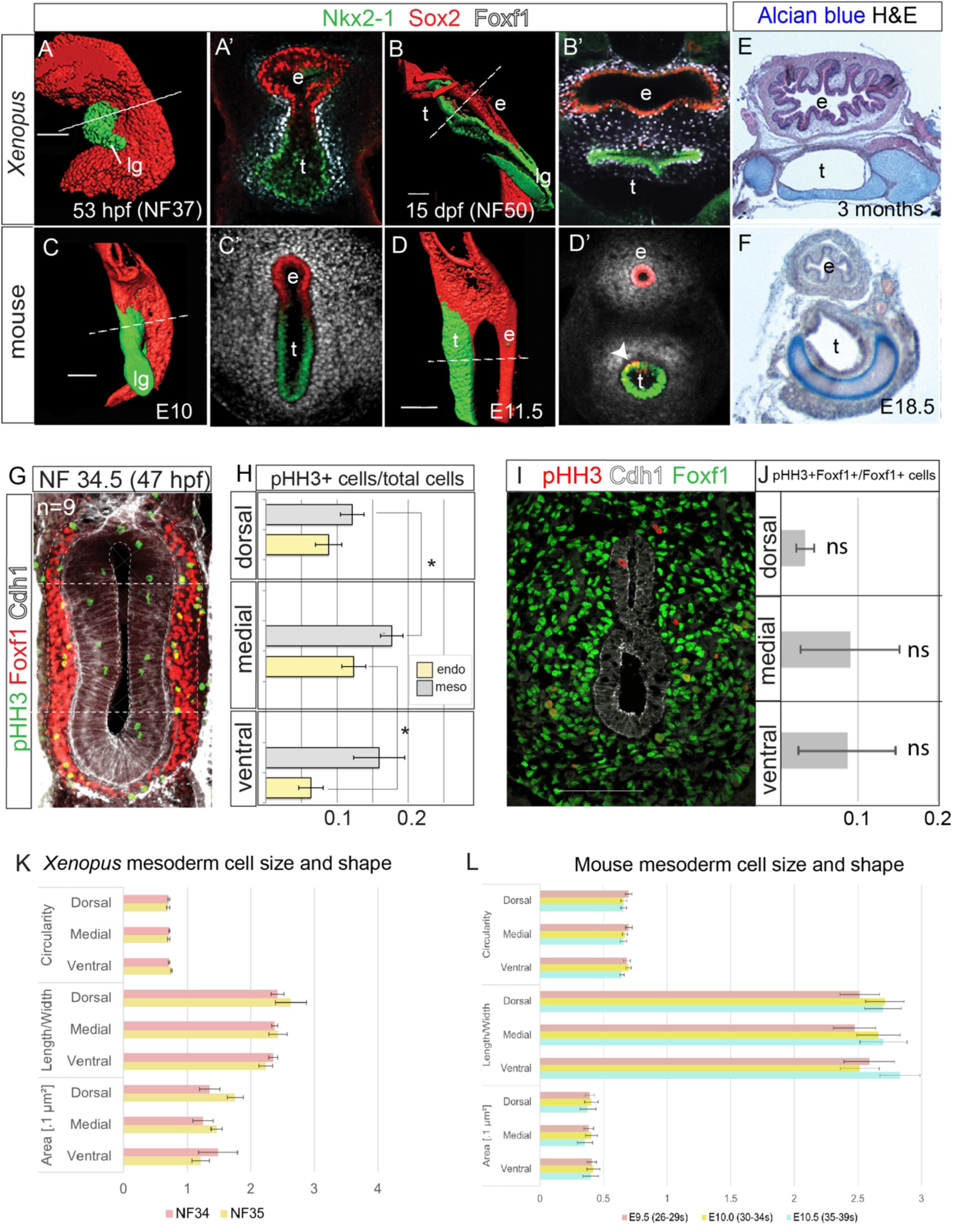
(Related to Figure 1) TE Morphogenesis is Conserved in *Xenopus* and Mouse. S1A-C: Wholemount confocal immunostaining of Sox2, Nkx2-1 and Foxf1 in *X. laevis* (A-B) and mouse (C-D) foregut epithelium. Surface renderings and optical sections show conserved TE morphogenesis, with Sox2+ dorsal foregut giving rise to esophageal epithelium and Nkx2-1+ central foregut giving rise to tracheal and lung epithelium. After separation, rare Sox2+/Nkx2-1+ cells are observed in the ventral esophagus and dorsal trachea (arrowhead in D’). S1E, F: Alcian blue cartilage staining shows similar tracheal differentiation in Xenopus (E) and mouse (F). e = esophagus, t = trachea, lg = laryngeal groove S1G: Immunostaining of phospho-histone H3-positive (pHH3+) cells, marking proliferating cells in NF34.5 *X. laevis*. S1H: Quantification of S1G. Endo = (pHH3+Cdh1+)/Cdh1+ cells, meso = (pHH3+Foxf1+)/Foxf1+ cells. The mitotic indices of the medial *X. laevis* endoderm and mesoderm are higher than those seen in the corresponding dorsal and ventral areas. Student’s two-tailed t-test, *p<0.05. S1I: Immunostaining of pHH3, Foxf1 and Cdh1 in mouse E10.0 foregut. S1J: Quantification of S1G, showing no statistically significant difference in pHH3+/Foxf1+ mesoderm cell proliferation from different regions. Two-way ANOVA, *p<0.05. S1K: Size and shape of *Xenopus* mesoderm cells. Average circularity, length/width ratio, and cell area (within .1µm^2^) of NF34 and NF35 foregut mesoderm show no statistical difference between regions. Mixed-effects analysis, *p<0.05. S1L: Size and shape of mouse mesoderm cells. Average circularity, length/width ratio, and cell area (within .1µm^2^) of E9.5, E10.0 and E10.5 foregut mesoderm show no statistical difference between regions. Mixed-effects analysis, *p<0.05.

**Supplemental Figure S2.**
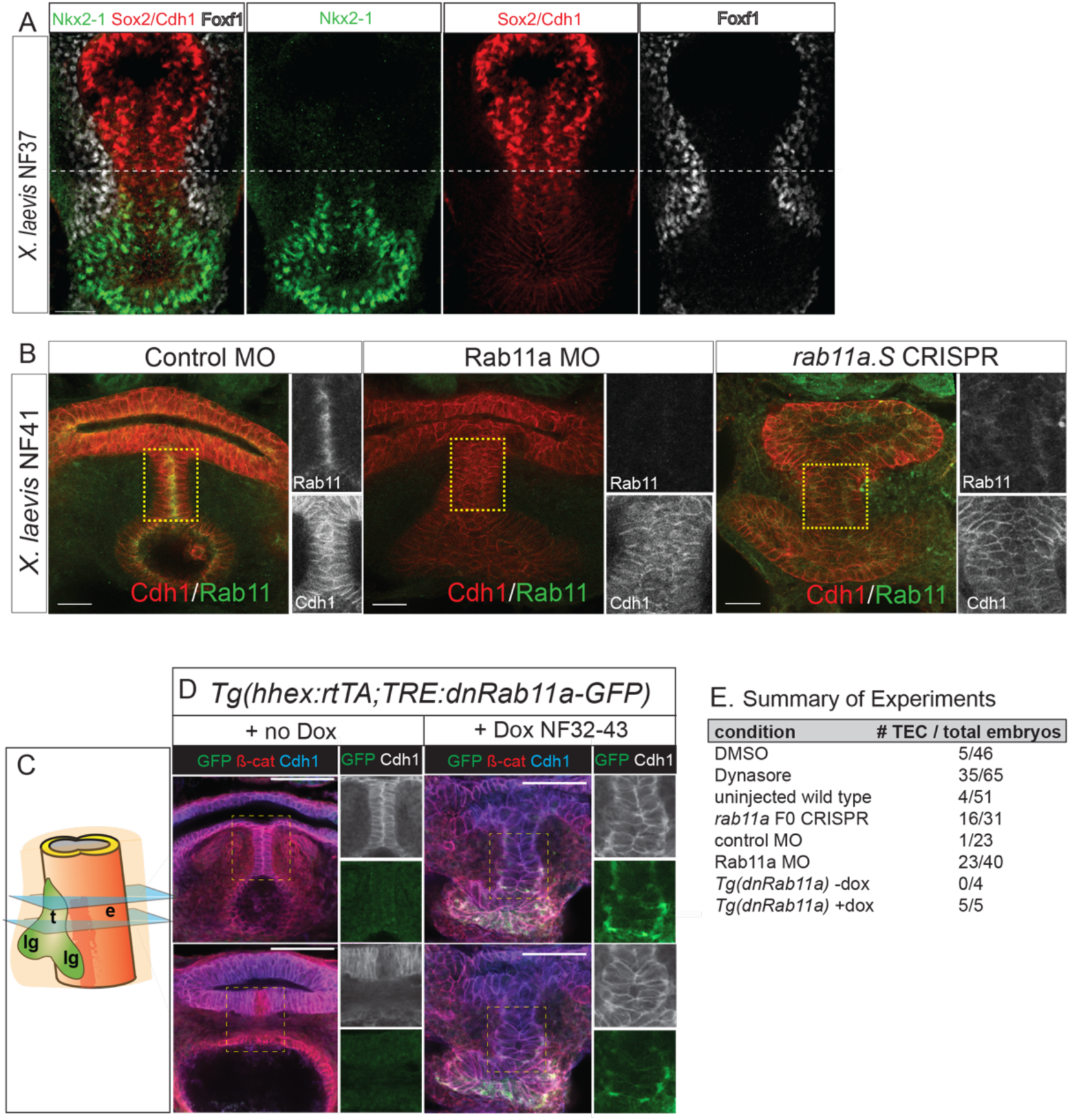
(Related to Figure 2) Rab11a-Dependent Epithelial Remodeling in the Transient Epithelial Septum that Co-Expresses Sox2 and Nkx2-1. S2A: Immunostaining of NF37 *X. laevis* embryos showing co-localization of Sox2 and Nkx2-1 in the transient epithelial septum where the foregut constricts, similar to mouse. S2B: Immunostaining of Control MO, Rab11 MO and *rab11.S* F0 CRISPR mutant *X. laevis* embryos at NF41 confirming reduced Rab11 protein levels and disorganized epithelium septum in Rab11 mutants relative to control embryos. Scale bar, 50 µm. S2C: Graphic indicating relative sections of the tracheoesophageal septum and separated trachea and esophagus seen in S2D. S2D: Sequential optical sections from wholemount immunostaining of transgenic *Tg(hhex:rtTA; TRE:dnRab11a-GFP) X. laevis* embryos at NF42-43. Addition of doxycycline (dox) from NF32-43 resulted in the expression of a dominant-negative Rab11a specifically in the *hhex+* foregut, which caused a failure of TE septation and a persistent TEC similar to Rab11 knockdown. GFP shows that the dnRab11a protein inappropriately localized to the basal-lateral surface compared to the localization of wildtype Rab11 at the apical surface and fusion point (Fig. 2SD). This confirms that Rab11 is specifically required for TE separation, independent of possible prior requirements for Rab11 during gastrulation. Scale bar, 100 µm. S2E: Summary of TEC frequencies in experiments that block Rab11-dependent endosome-mediated epithelial remodeling in *X. laevis* embryos.

**Supplemental Figure S3.**
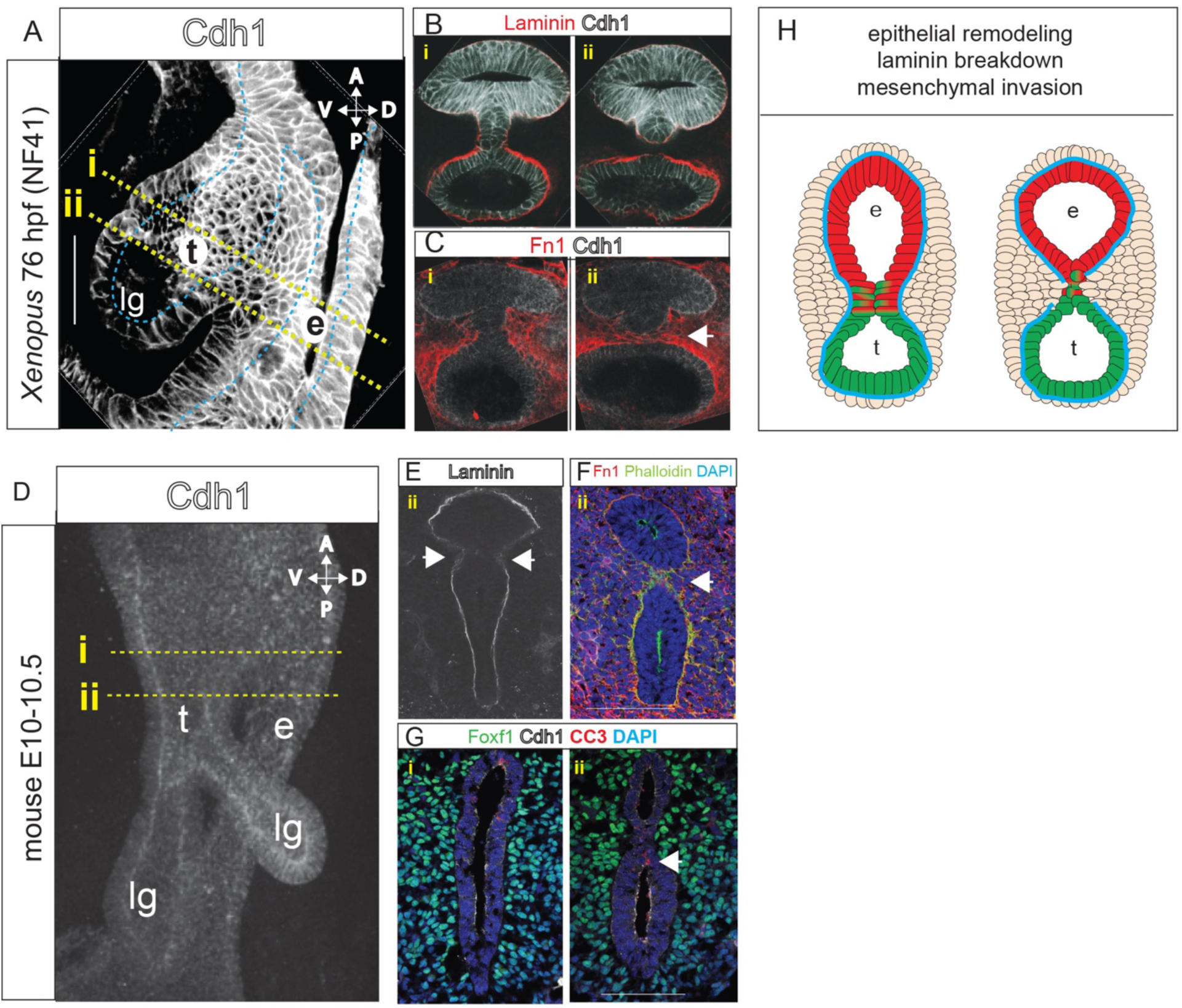
(Related to Figure 3) Resolution of the Epithelial Septum. S3A: Wholemount Cdh1 immunostaining of the NF41 *X. laevis* foregut (sagittal view) during (i) and immediately after (ii) TE septum resolution. Trachea (t) and esophagus (e). d; dorsal, v; ventral, a, anterior; p, posterior S3B: Optical section of Cdh1 and Laminin immunostaining in a NF41 *X. laevis* foregut during (i) and after (ii) TE septum resolution. Laminin decreases around the septum as the epithelial cells downregulate Cdh1 and transition from a columnar to round morphology. S3C: Optical section of Fibronectin (Fn1) and Cdh1 immunostaining in a NF41 *X. laevis* foregut during (i) and after (ii) TE septum resolution. Fn1-enriched mesenchymal cells are present in the midline between the nascent trachea and esophagus, suggesting cell migration. S3D: Wholemount Cdh1 immunostaining of an E10.5 mouse embryo (i: prior to TE separation, ii: during TE separation) showing the constriction of the foregut prior to TE separation. S3E: Immunostaining showing localized extracellular matrix degradation at the septation point (arrows) in an E11.0 mouse embryo. S3F: Immunostaining showing enrichment (arrow) of phalloidin and fibronectin just after TE separation in an E11.0 mouse embryo, suggesting medial mesenchymal cell movement. Scale bar, 100 µm. S3G: Cleaved caspase 3 (CC3)-positive cells, indicating apoptosis before (i) and during (ii) TE separation in an E10.5 mouse embryo. Presence of CC3+ cells in the endoderm during TE septum formation suggests that apoptosis may contribute to TE separation. Arrow denotes CC3 staining in the TE septum. Scale bar, 100 µm. S3H: Graphical summary of TE septum resolution, including epithelial remodeling at the midline fusion point, bilateral Laminin breakdown on either side of the septum, and mesenchymal invasion as the distinct trachea and esophagus form.

**Supplemental Figure S4.**
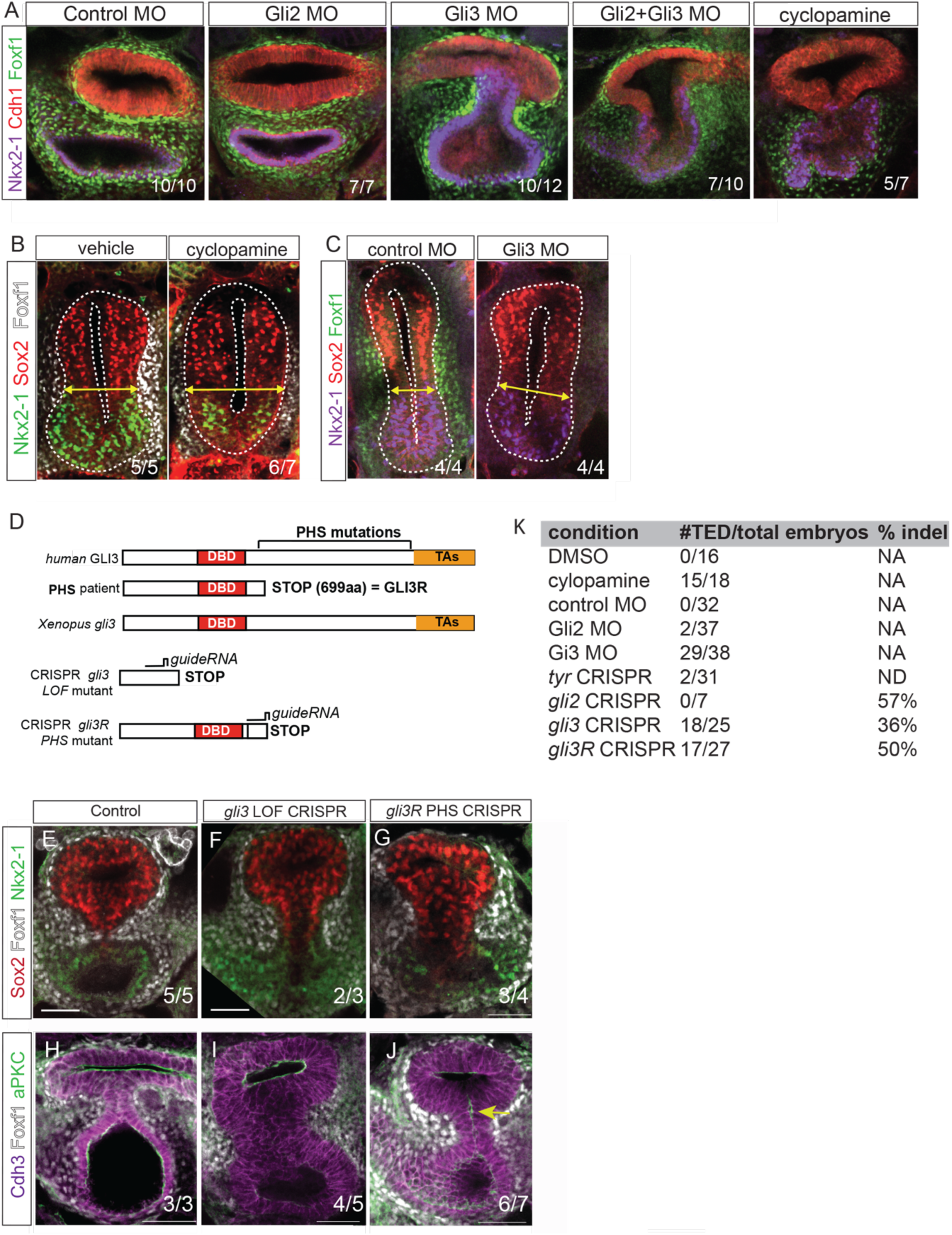
(Related to Figure 4) HH Signaling and Gli3 are Required for TE Morphogenesis in *Xenopus*. S4A: Immunostaining of Nkx2-1, Cdh1 and Foxf1 in NF42 *X. laevis* embryos injected with previously validated control MOs, Gli2 MOs *or* Gli3 MOs (5-7 ng) (Rankin et al 2016) or treated with the HH antagonist cyclopamine. Gli2 MO embryos show no defect, while Gli3 MO and Gli2+Gli3 MO fail to septate and exhibit a TE cleft. This suggests that *gli3* plays a more critical role than *gli2* during *X. laevis* TE separation. Note higher doses of Gli2+Gli3 MO (10 ng each) fail to pattern the foregut and induce Nkx2-1 as previously shown (Rankin et al 2016). S4B: Immunostaining of Nkx2-1, Sox2 and Foxf1 in NF35 *X. laevis* embryos treated with cyclopamine exhibit delayed medial constriction as indicated by the yellow arrows and reduced Foxf1+ mesenchyme, but preserve dorsal-ventral patterning, similar to *Shh-/-* mutant mice (Figure 4). S4C: Immunostaining of Nkx2-1, Sox2 and Foxf1 in NF35 *X. laevis* embryos injected with control MOs or Gli3 MOs, showing delayed medial constriction in Gli3 morphants. S4D: Schematic of human and Xenopus Gli3 proteins and the truncating mutation that results in Pallister Hall Syndrome (PHS). Summary of targeting strategy for CRISPR/Cas9-mediated indel mutagenesis of *X. tropicalis gli3* in embryos. Mutation of *gli3* exon 2 is predicted to result in loss of function (LOF) mutants whereas *gli3R* PHS mutants are predicted to maintain Gli3 transcriptional repressor activity. S4E-G. Immunostaining of Sox2, Foxf1 and Nkx2-1 in control (E) and CRISPR-mediated F0 mutant *X. tropicalis* embryos at NF41. *gli3* LOF mutants (F) and *gli3R* mutants (G) fail to septate and exhibit a persistent TEC, similar to Gli3 MO *X. laevis* embryos. S4H-J: Immunostaining of Cdh3, Foxf1 and aPKC in control (H) and CRISPR-mediated F0 mutant *X. tropicalis* embryos at NF41. *gli3* LOF mutants (I) show an abnormally thick TE septum that fails to resolve, whereas *gli3R* PHS mutants show persistent aPKC in the septum (arrow), similar to the *Foxg1Cre;Gli3T* mouse mutants, indicating that excessive Gli3R activity prevents remodeling of apical-basal epithelium polarity in the septum and prevents septation. S4K: Summary of the TED phenotypes in Xenopus embryos with disrupted HH/Gli3 activity. The F0 CRISPR/Cas9 mutagenesis results in embryos with mosaic mutations. The average percentage of damaging indel mutations from sequencing-TIDE genotyping (see methods) is shown.

## STAR METHODS

### CONTACT FOR REAGENT AND RESOURCE SHARING

Further information and requests for resources and reagents should be directed to and will be fulfilled by the lead contact, Aaron Zorn.

### EXPERIMENTAL MODEL AND SUBJECT DETAILS

*Xenopus* and all mouse lines described here were housed at Cincinnati Children’s Hospital Medical Center (CCHMC) and maintained according to the NIH Guidelines for the Care and Use of Laboratory Animals. All animals were housed in a 12-hour light-dark cycle with standard chow, and only healthy animals were used for experiments. Both mouse and *Xenopus* embryos were collected to analyze tracheoesophageal morphogenesis, and the sex of embryos was not recorded. All experiments were performed using guidelines approved by the CCHMC Institutional Animal Care and Use Committee (IACUC2019-0006 and IACUC2016-0059).

### METHOD DETAILS

#### Experimental Design

All *Xenopus* experiments were performed at least three times. In the case of mouse mutants, at least three mouse mutants, age-matched by somite number if collected at E11.5 or earlier, were used for analysis in each experiment. Sample size and animal sex were not deliberately determined. For most of the analysis samples were not blinded except the determination of TEC phenotypes in *Xenopus* embryos, which were scored blind. No data was excluded from analysis.

#### *Xenopus* experiments

*Xenopus laevis* and *Xenopus tropicalis* were purchased from Nasco or the National Xenopus Resource Center (*X. laevis* memGFP) and housed according to established CCHMC IACUC protocols. *X. tropicalis* “Superman” animals were generously provided by Mustafa Khokha (Yale University). Generation of *Xenopus* embryos was performed using ovulation and *in vitro* fertilization techniques previously described.

For *X.* laevis microsurgery experiments the ectoderm and lateral plate mesoderm surrounding the foregut were removed from one side of NF32/33 embryos and cultured in 0.5XMBS with gentamicin until NF35 when they were fixed along with unmanipulated siblings.

For morpholino (MO) knockdowns we injected previously validated Gli3-MOs (5 ng), Gli2-MO (5 ng) (Rankin et al., 2016) and Rab11a-MO (50 ng) (Kim et al., 2012), or equal concentrations of standard control morpholinos, into the dorsal-ventral region of 2-8 cell stage *X. laevis* embryos to target the future foregut region.

For transient F0 mosaic CRISPR/Cas9 mutagenesis, guide RNAs (gRNAs) were designed using CRISPRScan or CHOPCHOP and synthesized by IDT. For *gli2* and *gli3* loss of function, gRNAs targeting the *X. tropicalis gli2* exon 4 or *gli3* exon 2 were designed to produce truncating indels or missense mutations resulting in premature stop codon prior to the DNA-binding domains. A gRNA targeting the *X. tropicalis gli3* in exon13 corresponding to the human Pallister Hall Syndrome *GLI3*^*d699*^ mutation was designed such that a premature stop codon would prevent translation of the Gli3 activator domain resulting in a constitutive repressor form. For the *rab11a* LOF mutations, we designed gRNAs targeting a region in exon 2 that has identical sequence identity between and *X. tropicalis rab11a* and *X. laevis rab11a.S* (*X.laevis rab11a.L* has a 2-nucleotide mismatch and is predicted not to be targeted). *X. laevis* embryos were used for *rab11a* CRISPR/Cas9 mutagenesis because the Rab11 antibodies (to verify loss of Rab11) work in *laevis* but not *tropicalis*. RNA-seq data in Xenbase.org indicated that *rab11a.S* transcripts are expressed two-fold higher than *rab11a.L* in *X. laevis* tadpoles. Consistent with the mRNA levels, Rab11 immunofluorescence showed that *rab11a.S* CRISPR mutants (with wildtype *rab11a.L*), have dramatically reduced Rab11a protein levels (Figure S2B). To generate F0 mosaic *gli2, gli3, gli3R* and *rab11a.S* mutant embryos, the gRNAs (750 pg) were then injected with Cas9 protein (1-1.5 ng; PNA Biosciences) into *X. tropicalis* or *X. laevis* embryos at the one-cell stage or the eight-cell stage targeted to the foregut endoderm. For negative controls, *tyrosinase* (*tyr*) gRNAs were injected. These resulted in a high frequency of F0 embryos lacking pigment indicating effective indel mutations, but no detectable TED.

To genotype embryos, genomic DNA was extracted from tails and the genomic regions target by the gRNAs were PCR amplified with 2X Phusion Master Mix (Thermo Fisher Scientific F531) according to manufacturer’s instructions. Successful indel mutation was initially verified with a T7E1 (New England BioLabs Incorporated M0302S) digest of the PCR product, followed by, Sanger sequencing. Overall efficiency and allele frequency of specific mutations were determined by TIDE sequencing decomposition analyses (https://tide.deskgen.com). The table below shows the efficiency of truncating indel mutations for each gRNA.

| Gene, exon (species) | Damaging indel %, n |
| --- | --- |
| *gli2* exon 4 (*X. tropicalis*) | 57% (n=7) |
| *gli3* exon 2 (*X. tropicalis*) | 36% (n=25) |
| *gli3* exon 13 PHS (*X. tropicalis*) | 50% (n=27) |
| *rab11a.S* exon 2 (*X. laevis*) | 78% (n=16) |

For small molecule treatments, *Xenopus laevis* embryos were cultured at room temperature, covered in aluminum foil, in 0.1XMBS in the presence of small molecules that were refreshed once every 12-24 hours beginning at NF28/29-32/33. Control sibling embryos were treated with equal volumes of the small molecule solvent (e.g., EtOH or DMSO). Small molecule treatment concentrations were: cyclopamine (Selleck or Tocris, 80µM), Dynasore (Millipore Sigma, 50µM), GM6001 (Millipore Sigma, 150µM) and 1,10-phenanthroline (Millipore Sigma, 5 µM).

For Xenopus transgenesis, the I-SceI meganuclease method was used as described (Sterner et al., 2019). The *Tg(hhex;rtTA;TRE:dnRab11a-GFP)* construct was generated as follows. The −1.2Kb *X.laevis hhex.L* promoter (Rankin et al., 2011) and the dNRab11a-GFP fusion coding sequence (Kim et al., 2012) were PCR amplified gel purified and TOPO-TA cloned into the pCR8 Gateway entry vectors (Thermo Scientific #K250020). Gateway LR Clonase II Plus enzyme (Thermo Scientific #12538120) was used in standard recombination reactions according to manufacturer’s instructions to transfer the *hhex.L* promoter into the pDXTP and dNRab11-GFP into the pDXTR transgenesis plasmids (Sterner et al., 2019).

Transgenic embryos were generated as follows: 200pg of pDXTP-*hhex* promoter and 200pg of pDXTR-dNRab11-GFP were incubate in a 25uL reaction containing 2.5uL of I-SceI meganuclease enzyme (New England Biolabs #R0694S) in 0.5X I-SceI buffer. The reaction was incubated at 37°C for 30 to 40 minutes and then immediately injected into 1-cell embryos near the sperm entry point within the first 45 to 60 minutes after fertilization. Embryos were cultured at 13°C for the first two hours after injection and subsequently at 18°C to 23°C degrees thereafter. Transgenic embryos were selected based on GFP fluorescence in the eye, which becomes visible during early tailbud stages (Sterner et al., 2019). In half of the embryos the transgenes were activated by the addition of Doxycycline hyclate (Sigma #D9891) at a final concentration of 50ug/mL culture buffer. The addition of Dox alone has no effect on the embryos and did not cause TEDs.

All of the confocal analysis in *Xenopus* embryos was performed in wholemount. Embryos were cut transversely just anterior to the pharynx and posterior to the lung buds in 100% MeOH. After serial rehydrations into 1XPBS with 0.1% Triton X-100 (1XPBSTr), embryos underwent antigen retrieval for 45 minutes in 1X citrate buffer at 65°C. Following three washes in 1XPBSTr, embryos were blocked in a 1XPBSTr solution of 1% bovine serum albumin (BSA) and 5% DMSO overnight at 4°C on a rocker for at least sixteen hours. Primary antibodies were added to embryos for at least sixteen hours, again rocking overnight at 4°C. Following five thirty-minute washes in 1XPBSTr, secondary antibodies were added in a 1XPBSTr solution containing 0.1% BSA overnight, rocking at 4°C for at least sixteen hours. After five thirty-minute washes in 1XPBSTr, embryos were serially dehydrated into 100% methanol, and stored at 4°C until imaging in Murray’s Clear solution.

#### Mouse experiments

All mouse experiments performed were approved by the CCHMC Institutional Animal Care and Use Committee (IACUC). *Gli2*^*tm2.1Alj*^*/J* mice were obtained from The Jackson Laboratory. *Foxg1*^*Cre*^ and *Shh*^*CreGFP*^ mice were obtained from Debora Sinner’s lab. *Gli3T*^*Flag*^ mice were provided by Rolf Stottmann and Joo-Seop Park. *Gli3*^*XtJ*^ mice were obtained from Rolf Stottmann’s lab. *Nkx2-1*^*GFP*^ mice were provided by Jeffrey Whitsett and John Shannon. *FoxA2*^*tm2.1(cre/Esr*)Moon/J*^ and *Sox2*^*tm1.1Lan/J*^ mice were provided by James Wells (tamoxifen administered via oral gavage at E6.5). Mice were maintained in the CCHMC animal facility. Timed matings and somite counting were used to obtain embryos at the stages described. Genotyping was performed using Phusion Hot Start Flex 2x Master Mix or Quickload 2x Master Mix (New England BioLabs, Incorporated). After collection, all mouse embryos were washed in 4% paraformaldehyde, rocking overnight at 4°C, before two five-minute washes in 1XPBS.

Mouse embryos used for cryosectioning were washed for one to two days, rocking at 4°C, in 30% sucrose in 1XPBS. Embryos were then embedded in OCT (Sakura, VWR) and stored at −80°C until sectioning at 8 µm thickness or 60 µm thickness. For section immunohistochemistry, sections were washed in 1XPBS, then 1XPBS with 0.1% Triton X-100 for permeabilization before blocking for one hour in 5% normal donkey serum in 1XPBS. After incubation in primary antibodies in 1XPBS at 4°C overnight, slides were washed in 1XPBS before incubation in secondary antibodies in 1XPBS at room temperature for one hour, followed by 1XPBS washes and mounting using ProLong Gold Antifade (Thermo Fisher Scientific).

Mouse embryos used for wholemount immunostaining were incubated in 100% methanol at −20°C until dissection and staining of the foregut. Briefly, embryos were washed for two hours at room temperature in Dent’s Bleach solution before serial rehydration into 1XPBS from methanol. Embryos were blocked in 5% normal donkey serum in 1XPBS with 0.2% Triton X-100 for two hours, rocking at room temperature. Embryos were then placed in primary antibodies in blocking solution, rocking overnight at 4°C. After a series of washes in 1XPBS with 0.1% Triton X-100, embryos were incubated with secondary antibodies in blocking solution, rocking overnight at 4°C. After three washes in 1XPBS with 0.1% Triton X-100, embryos underwent serial dehydration into 100% methanol before overnight storage at 4°C. Embryos were removed from methanol and placed into Murray’s Clear for at least fifteen minutes before confocal imaging.

### QUANTIFICATION AND STATISTICAL ANALYSIS

NIS Elements software was used to obtain image quantifications. For cell proliferation and cell death counts, the number of pHH3+ or CC3+/Cdh1+ and pHH3+ or CC3+/Foxf1+ cells were counted, as well as pHH3+ or CC3+/DAPI+ cells when possible. In *Xenopus* and mouse the mesoderm thickness in the dorsal, medial or ventral regions was measured as the distance between the lateral ectoderm and endoderm. Mesoderm density in *Xenopus* was calculated as the number of Foxf1+ cells within a confocal Z-stack ∼10^5^ µm^3^, whereas the cell density in mouse was calculated based on DAPI staining from individual 60µm confocal optical sections.

For small molecule treatments, the laryngotracheal groove was calculated as the distance between the start of the groove and the distinct trachea, marked by absent cytoskeletal staining between the trachea and esophagus. The tracheal length was calculated as the distance between the start of the distinct trachea, described above, and the start of the lung buds. Data shown is from one representative experiment. Pairwise significance calculations were performed in Microsoft Excel using a Student’s two-tailed t-test. All other significance calculations were performed using one-way ANOVA (for one control with multiple conditions), two-way ANOVA (for two controls with equal numbers of conditions), or mixed effects analysis (for two controls with unequal numbers of conditions) in Prism. p < 0.05 indicated significance for all tests, and the average and SEM were included in all calculations.

## KEY RESOURCES TABLE

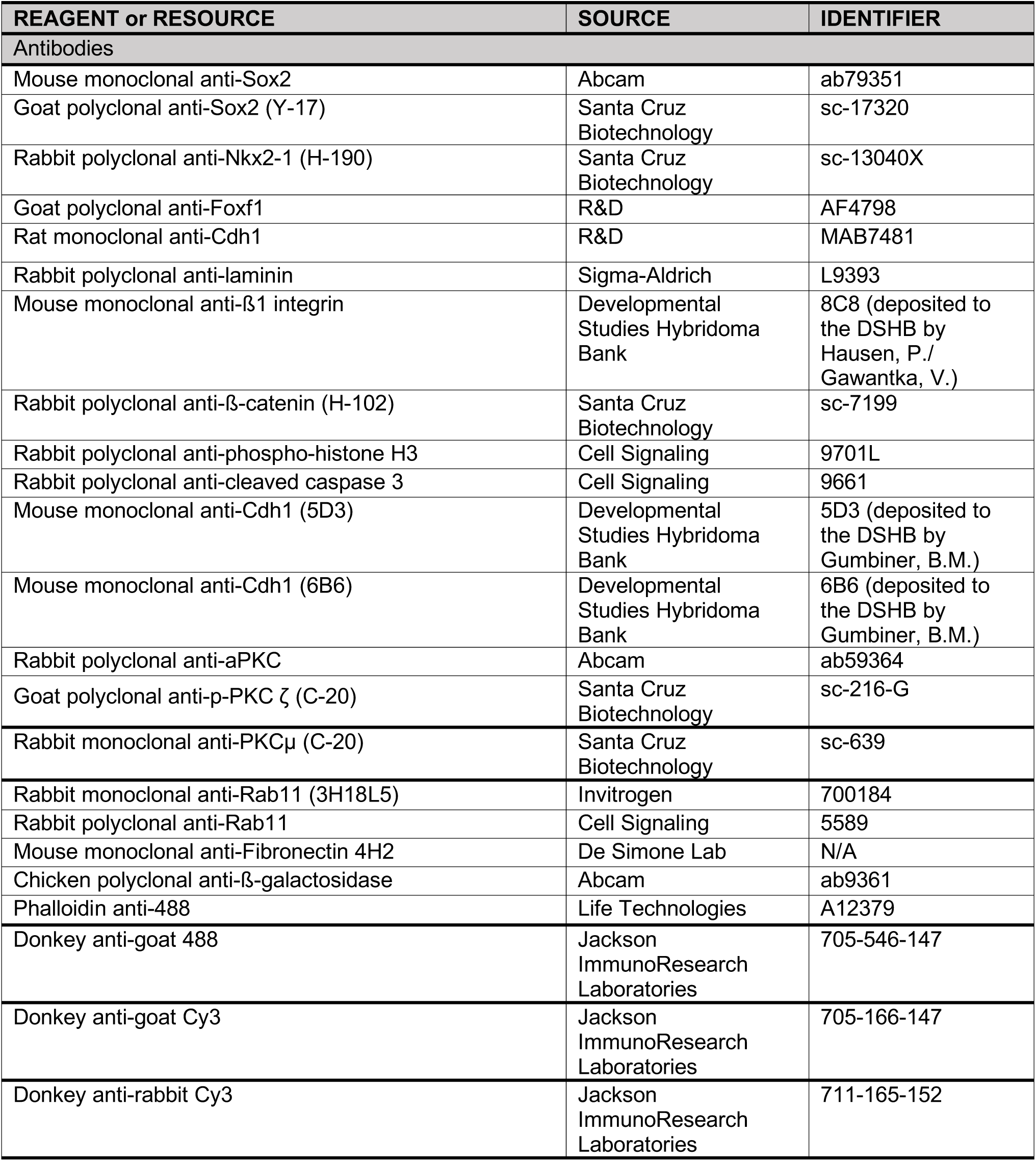
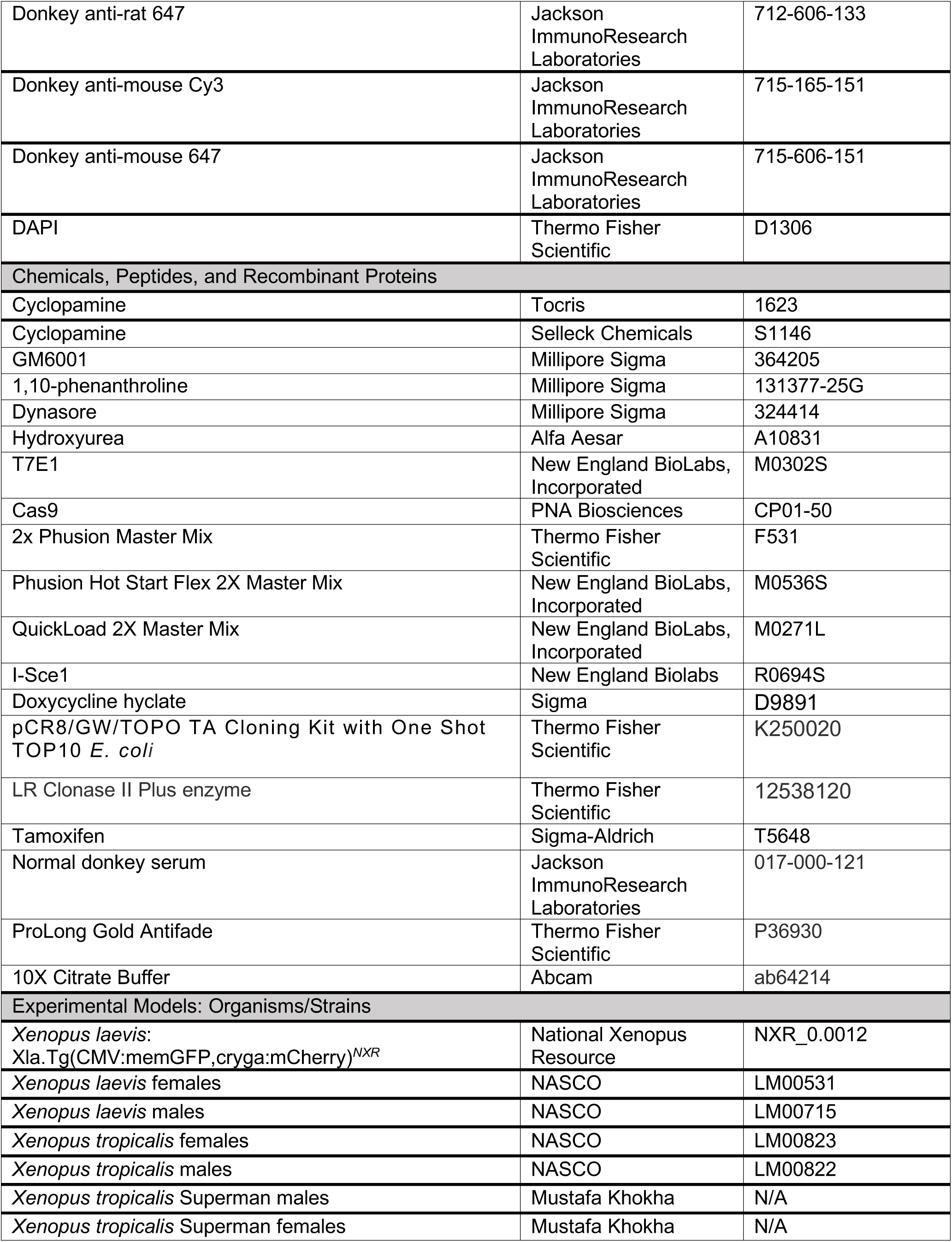
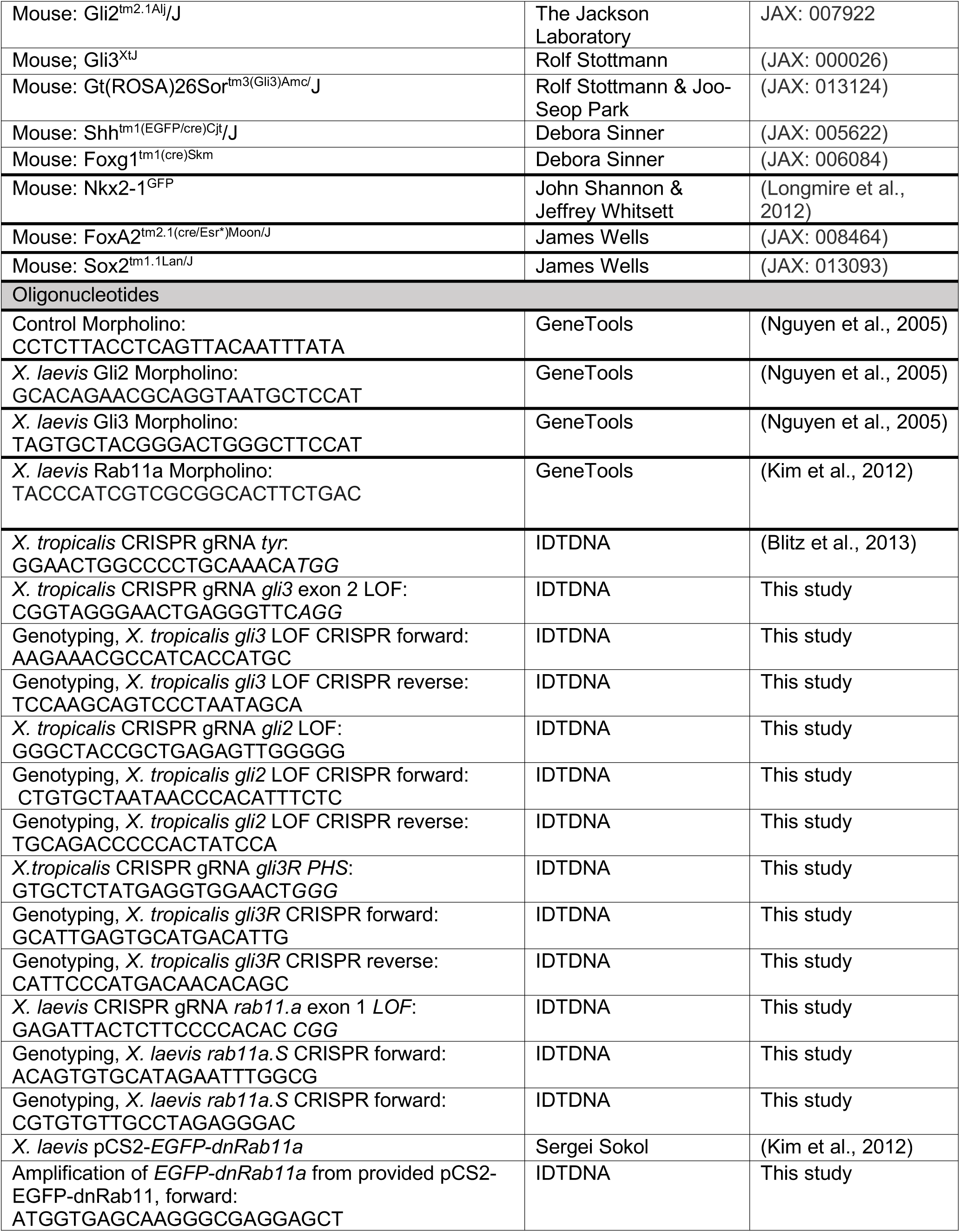
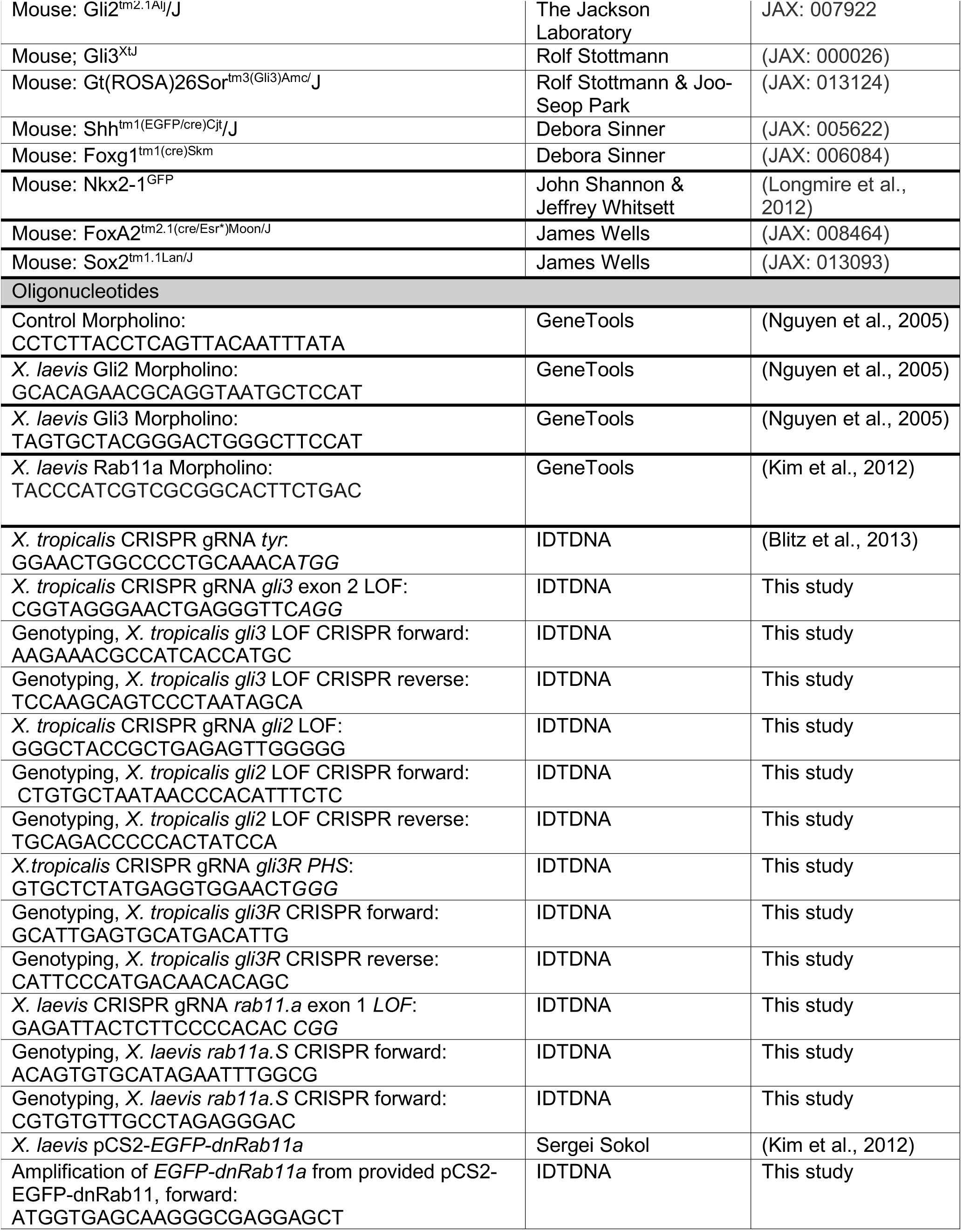
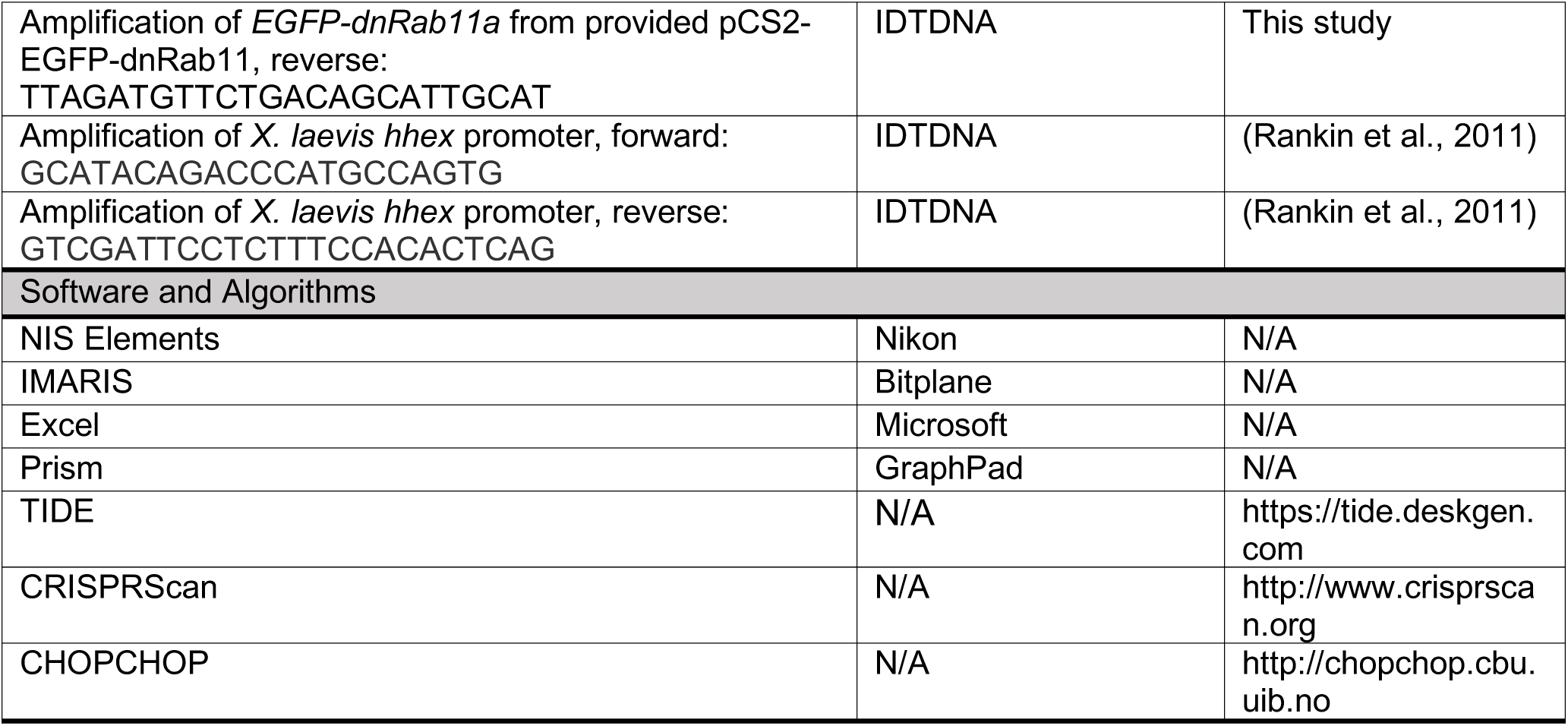

